# The Herbicide Glyphosate Promotes Hypertension via Gut Microbiota-Mediated Mechanisms

**DOI:** 10.64898/2026.02.25.708064

**Authors:** Ishan Manandhar, Sudhan Pachhain, Ramakumar Tummala, Blair Mell, Arturo Grano De Oro, Sachin Aryal, Xue Mei, Maya Nair, Sanjana Kumariya, Wisdom Ahildja, Oluwatosin Mautin Akinola, Pritam Bardhan, Tao Yang, Beng San Yeoh, Yuan Tian, Andrew D. Patterson, Zhong-min Li, Kurunthachalam Kannan, Matam Vijay-Kumar, Islam Osman, Piu Saha, Bina Joe

## Abstract

Glyphosate, the active ingredient in herbicide Roundup, is the most widely used environmental contaminant that has been extensively studied for its potential carcinogenic effects. In the US alone, a staggering 81% of the US population ≥6 years of age is exposed to glyphosate. Notably, this coincides with the alarming rise in the incidence of hypertension, the single largest risk factor for global mortality through cardiovascular diseases. Here we asked if there is a link between glyphosate exposure and hypertension, the premise being that glyphosate targets the shikimate pathway present in gut microbiota coupled with more recent knowledge that gut microbiota causally regulate blood pressure. We hypothesized that glyphosate elevates hypertension through microbiota-mediated mechanisms and document a highly concerning detrimental effect of glyphosate by demonstrating a causal link between oral glyphosate exposure and significant elevation in blood pressure. Gut microbiota was identified as the central mediator of this effect. Mechanistically, glyphosate-mediated elevation in blood pressure was through disruption of both gut–liver and gut-vascular homeostasis via FXR-signaling and accumulation of the microbial metabolite shikimic acid, respectively. Together, these findings underscore the need to reconsider the unabated use of this herbicide, which adversely affects a cardinal sign of health.

## Introduction

In the late 1970s, Monsanto introduced a highly effective herbicide and marketed it as Roundup^1^. Since then, the active ingredient of Roundup, glyphosate has remained the most widely used broad-spectrum herbicide in the history of agriculture^2,3^. It is used worldwide in over 750 different products for agriculture, forestry, urban, and home garden applications^4^. Human exposure to glyphosate can occur through dermal contact, inhalation, and diet^5^. Glyphosate exposure has been documented in several occupational and population studies worldwide. More recently, an analysis of glyphosate from urine samples obtained from the US general population from the 2013-2014 National Health and Nutrition Examination Survey (NHANES) revealed that glyphosate was detected in a staggering 81% of the US population ≥6 years of age^6^. Intriguingly, data from the NHANES study also shows that the incidence of hypertension, the leading risk factor for cardiovascular diseases, is also significantly on the rise^7^.

The estimated environmental risk for hypertension outweighs its genetic risk, leading us to consider yet uninvestigated environmental factors driving hypertension. Glyphosate stood out as one such candidate because of its mode of action. Glyphosate inhibits the enzyme, 5-enolpyruvyl shikimate-3-phosphate synthase (EPSPS), which is exclusively present in plants, but not in mammals^8^. However, microbes also possess EPSPS^9,10^. Recent recognition that humans are an ecosystem consisting of trillions of microbes residing within the human gastrointestinal tract^11,12^ led us to consider the effects of glyphosate on gut microbiota as a cause for concern. This is because we and others have shown that alterations in gut microbiota composition causally impact blood pressure^13–15^. Using genetically hypertensive rats, we have previously demonstrated that treatment with antibiotics to eliminate gut microbiota increased blood pressure^16^. Similar modulations of gut microbiota using a variety of methods have been demonstrated to elevate blood pressure^17–20^.

Based on these observations, in the current study we hypothesized that glyphosate elevates blood pressure. To test this hypothesis, we used experimental rats, with a genetic predisposition to develop hypertension^21^ and exposed these rats to glyphosate by oral administration. Our results illustrate that glyphosate is a pro-hypertensive environmental hazard.

## Results

### Exposure to Glyphosate Elevated Blood Pressure

Two groups of male Dahl Salt-Sensitive (S) rats were implanted with radiotelemetry transmitters to monitor blood pressure and exposed to 175 mg/kg BW/day of glyphosate in their drinking water *ad libitum*. Compared to controls without this exposure, a highly significant elevation in mean arterial pressure (MAP) (148 mmHg vs. 142 mmHg, p<0.0001) and systolic blood pressure (173 mmHg vs. 162 mmHg, p<0.0001) were observed (**Supplementary Figures 1A-C**) after one week of exposure. By the third week of glyphosate exposure, the differences were more pronounced, with notable elevation of MAP (168 mmHg vs. 155 mmHg, p<0.0001), systolic blood pressure (195 mmHg vs. 176 mmHg, p<0.0001) and diastolic blood pressure (142 mmHg vs. 135 mmHg, p<0.0001) (**Figure 1A-C).**

**Figure 1.**
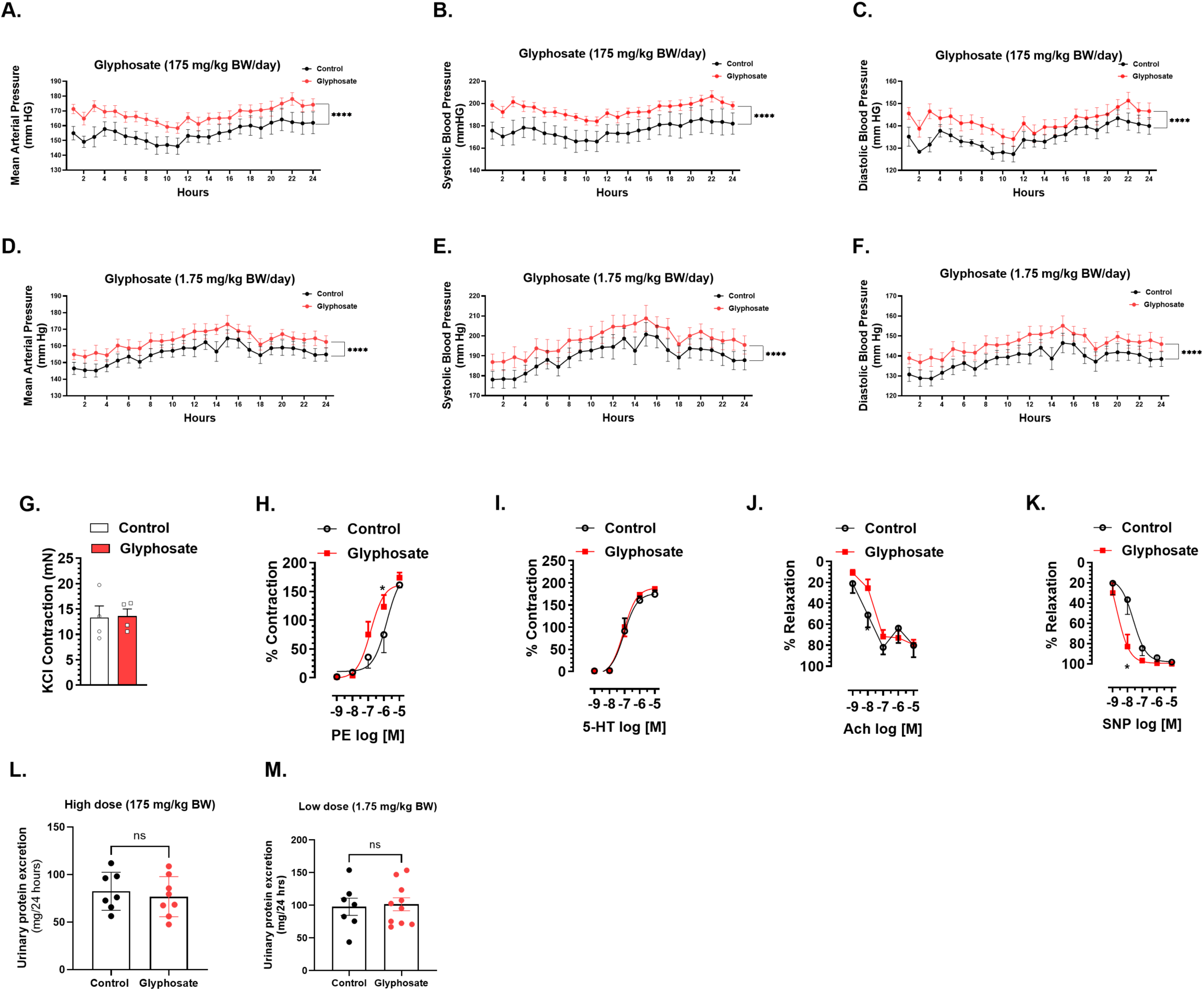
Glyphosate Administration Increases Blood Pressure in Dahl Salt-Sensitive (S) rats. **(A–C)** Blood pressure comparisons between control and glyphosate-treated rats (175 mg/kg body weight daily): **(A)** Mean arterial pressure **(B)** Systolic blood pressure **(C)** Diastolic blood pressure (n = 7–8/group). **(D–F)** Blood pressure comparisons between control and glyphosate-treated rats (1.75 mg/kg body weight daily): **(D)** Mean arterial pressure. **(E)** Systolic blood pressure **(F)** Diastolic blood pressure rats (n = 7–8/group). **(G–K)** Vascular reactivity analysis in mesenteric resistance arteries (n = 4/group): **(G)** Contraction induced by KCl; **(H)** Contractions induced by phenylephrine (PE, 10⁻□–10⁻□M); **(I)** Contractions induced by serotonin (5-HT, 10⁻□–10⁻□M); **(J)** endothelium-dependent relaxations induced by acetylcholine (ACh, 10⁻□–10⁻□M); **(K)** endothelium-independent relaxations induced by sodium nitroprusside (SNP). **(L–M)** Urinary protein excretion comparisons between glyphosate-treated and control rats: **(L)** 175 mg/kg BW/day; **(M)** 1.75 mg/kg BW/day. Data are presented as mean ± SEM. * p < 0.05, ** p < 0.01, **** p < 0.0001. Data are presented as mean ± SEM. ns = non-significant, * p < 0.05, **** p < 0.0001.

Consistent with these blood pressure changes, *ex vivo* wire myography of isolated mesenteric resistance arteries from rats exposed to glyphosate revealed significantly increased sensitivity to phenylephrine (PE) and a decreased sensitivity to the vasorelaxation responses to increasing concentrations of acetylcholine (Ach) (**Figures 1G-K**), indicating impaired vascular function. However, urinary protein excretion was comparable between the groups (**Figures 1L, M**).

For translational relevance, we further tested the effect of low-dose (1.75 mg/kg BW/day) glyphosate exposure on blood pressure. Here again, within a span of one week, rats exposed to glyphosate demonstrated a significant increase in MAP (150.1 mmHg vs. 146.0 mmHg, p<0.001), systolic blood pressure (182.2 mmHg vs. 177.8 mmHg, p<0.01) and diastolic blood pressure (134.1 mmHg vs. 130.1 mmHg, p<0.001) (**Supplementary Figures 1D-F**). This elevation in blood pressure at 4 weeks was sustained with a greater increase in MAP (163.2 mmHg vs. 155.4 mmHg, p<0.0001), systolic blood pressure (197.8 mmHg vs. 189.8 mmHg, p<0.0001) and diastolic blood pressure (145.9 mmHg vs. 138.1 mmHg, p<0.0001) (**Figures 1D-F**).

### Glyphosate Exposure Significantly Remodeled Gut Microbiota Composition

To examine whether oral exposure to glyphosate increased circulating glyphosate, its levels were quantified in both serum and urine. Data presented in **Figures 2A, B** confirmed that glyphosate was elevated both in circulation and excreted in high amounts. Next, fecal microbiota was profiled between groups of rats administered with and without glyphosate. Both 16S rRNA gene and whole genome sequencing approaches revealed significantly lower alpha diversity (Shannon diversity) in glyphosate-exposed rats (**Figure 2F**). Additionally, principal coordinate analysis (Bray-Curtis, beta diversity) showed distinct separation between the control and treated groups, demonstrating that oral glyphosate exposure significantly remodeled gut microbiota composition (**Figure 2G**). Sequencing by the 16S approach showed shifts in specific bacterial genera, with prominent decreases in abundances of *Parasutterella*, *Blautia* and *Streptococcus* and increasing abundances *of Erysipelatoclostridium*, *Ruminococcus* and *Lactobacillus* in glyphosate-exposed rats compared to controls (**Supplementary Figure 1G**). Similarly, whole genome sequencing further identified bacterial taxa at the species level. Notably, *Lactobacillus johnsonii*, *Bifidobacterium pseudolongum*, *Faecalibacterium rodentium*, and *Phocaeicola vulgatus* were decreased, whereas *Ligilactobacillus murinus*, *Ligilactobacillus animalis*, and *Clostridium sporogenes* were enriched in the group exposed to glyphosate (**Figure 2H**).

**Figure 2.**
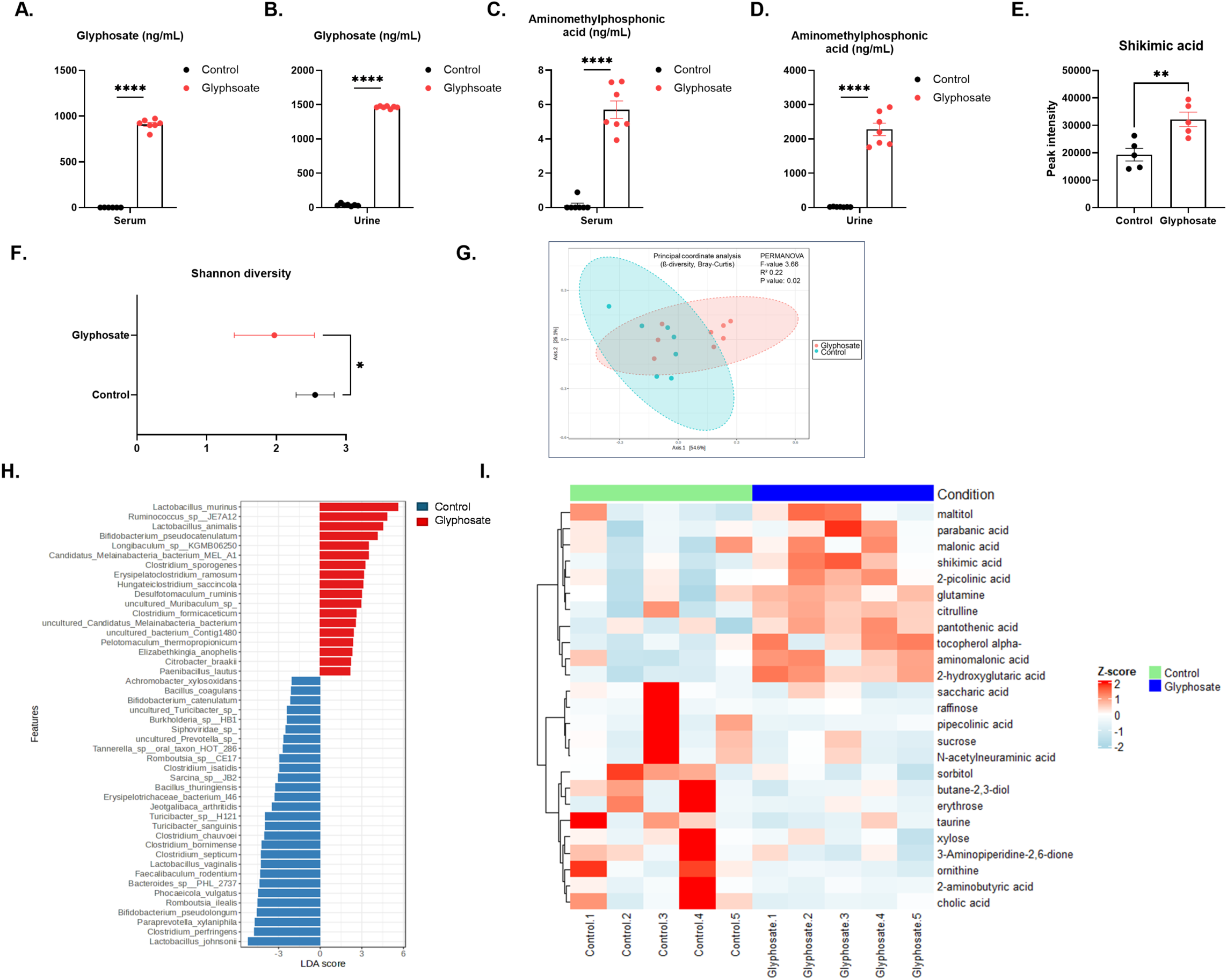
Glyphosate Exposure Remodels Gut Microbiota and Alters Serum Metabolites. **(A, B)** Comparisons of glyphosate levels in the serum and urine samples, **(C, D)** Comparisons of aminomethylphosphonic acid levels in the serum and urine samples, **(E)** Comparison of Shikimic acid levels in the serum samples between control and glyphosate-treated rats (175 mg/kg body weight daily). n=5-8/group. **(F)** Shannon diversity index of the gut microbiota in control and glyphosate-treated rats. **(G)** Principal Coordinate Analysis Plot showing significant differences in beta-diversity of fecal microbiome of the control and glyphosate-treated rats as analyzed by PERMONOVA (F-value=3.66, R^2^=0.22, p-value<0.05). n=7-8/group. **(H)** Gut bacterial whole-genome sequencing (by Oxford Nanopore GridIon) shows significant alterations at the species levels in glyphosate-treated S rats. Bar plots in red: elevated abundances in glyphosate-treated rats; bar plots in blue: diminished abundances in glyphosate-treated rats. Data was analyzed by linear discriminant analysis effect size (LEfSE). LDA≥2. **(I)** Heatmap of representative mass spectrometry peak intensities of serum metabolites in control and glyphosate-treated rats. Data are presented as mean ± SEM. *p < 0.05, **p < 0.01, ****p < 0.0001.

### Obligatory Role of Gut Microbiota as a Mediator of the Adverse Effect of Glyphosate on Blood Pressure

To determine the importance of gut microbiota in mediating the detrimental effect of oral glyphosate on blood pressure, we next administered glyphosate subcutaneously to bypass the gastrointestinal tract and avoid exposure to gut microbiota. In this study, blood pressure remained comparable between glyphosate-exposed and control rats (**Figure 3A**). Further, vascular function data collected via *ex vivo* wire myography using isolated mesenteric arteries were also comparable between subcutaneous glyphosate-exposed and control rats (**Figure 3B-F**). Together, these findings support the hypothesis that gut microbiota is an underlying, critical mediator of the pro-hypertensive effect of glyphosate.

**Figure 3:**
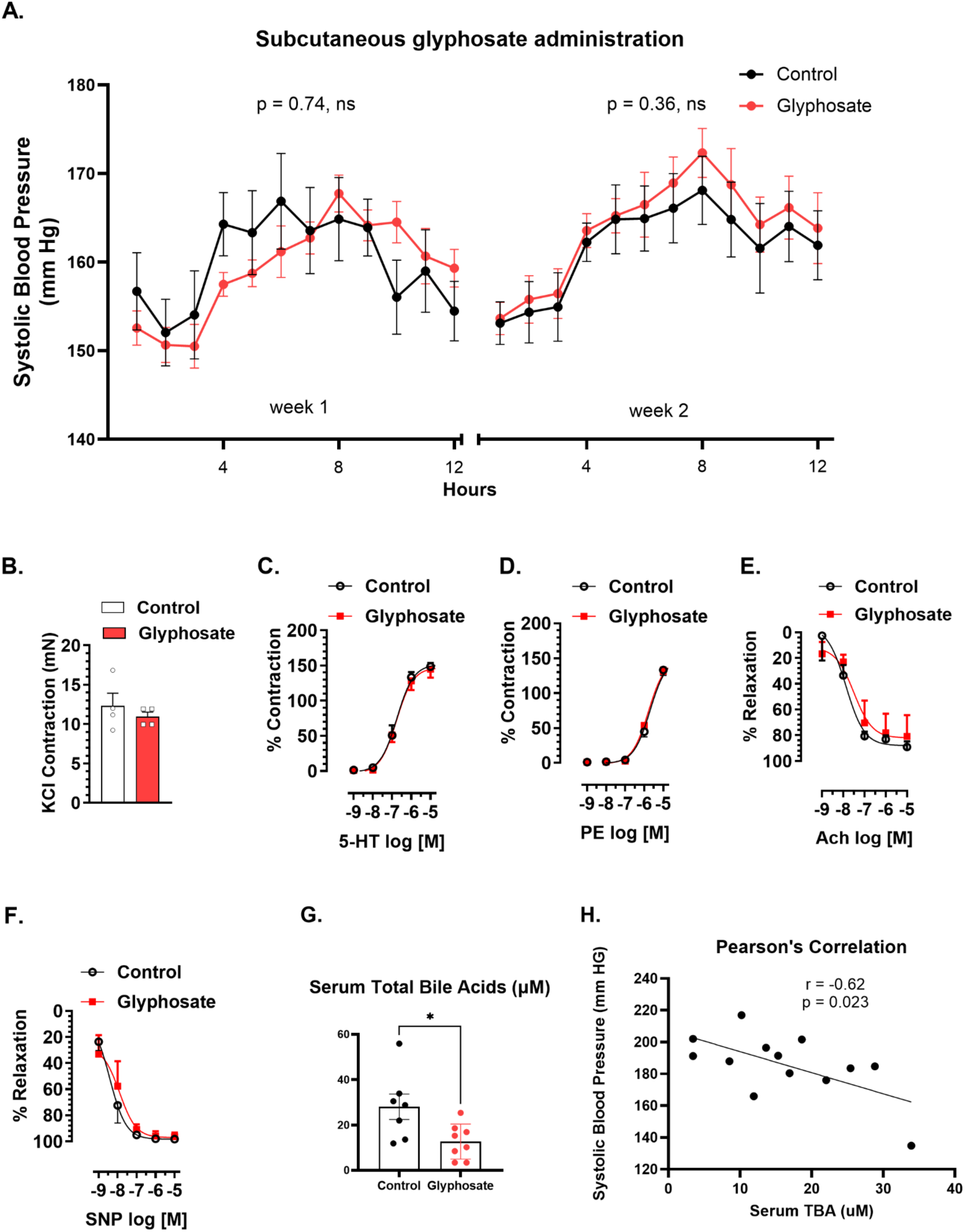
Subcutaneous Glyphosate Administration does not Increase Blood Pressure nor Alter Vascular Function. **(A)** Systolic blood pressure comparisons between control (n = 5) and glyphosate-treated (n = 5) (subcutaneous, 12 mg/day) male S rats at week 1 (160 mmHg vs 159 mmHg) and week 2 (162 mmHg vs 164 mmHg), ns; not statistically significant. **(B)** KCl-induced contraction in mesenteric resistance arteries from control and glyphosate-treated rats. **(C)** Contractions induced by serotonin (5-HT, 10⁻□–10⁻□M). **(D)** Contractions induced by phenylephrine (PE, 10⁻□–10⁻□M). **(E)** Endothelium-dependent relaxations induced by acetylcholine (ACh, 10⁻□–10⁻□M). **(F)** Endothelium-independent relaxations induced by sodium nitroprusside (SNP, 10⁻□– 10⁻□M). Data are presented as mean ± SEM. ns = non-significant. **(G)** Serum total bile acid levels in glyphosate-treated and control rats (n =7-8/group). **(H)** Correlation analyses between serum TBA and systolic blood pressure in rats treated with and without glyphosate (175 mg/kg BW). Systolic blood pressure is inversely correlated with the serum TBA. Pearson’s correlation, r =-0.62, p<05

### Glyphosate Exposure Significantly Remodeled Bile Acid Composition

Next, we tested whether the remodeled microbiota had any effect on the composition of bile acids, which are metabolites generated collaboratively by microbiota and the host. Serum total bile acids (TBA) were significantly lower in the rats exposed to glyphosate compared to controls (**Figure 3G**). Targeted bile acid profiling further confirmed that several classes of bile acids were lowered in the glyphosate-exposed group (**Supplementary Figure 2**). Interestingly, lower levels of serum total bile acids were negatively correlated with systolic blood pressure (r=-0.62, p=0.02) (**Figure 3H**) as confirmed by Pearson’s correlation. These data suggested that host signaling via bile acids could mechanistically contribute to glyphosate-induced hypertension.

### Farnesoid X receptor (FXR) Signaling Mediates Glyphosate-induced Hypertension

Since bile acids signal majorly via FXR, which is a key negative regulator of bile acid synthesis, we examined whether disruption of FXR signaling contributes to the acute elevation of blood pressure caused by glyphosate. Using the CRISPR/Cas9-mediated genome editing technology, targeted disruption of the locus encoding FXR*, Nr1h4, was* achieved in the genetic background of Dahl Salt-Sensitive (S) rats (**Figures 4A-C**). Consistent with previous reports, ablation of FXR resulted in a significant elevation of total serum bile acids in *Fxr* knockout (*Fxr*KO) rats compared to S rats (**Figure 4D**). As expected, several classes of bile acid levels were elevated in *Fxr*KO rats as shown in **Supplementary Figures 3A-C**.

**Figure 4:**
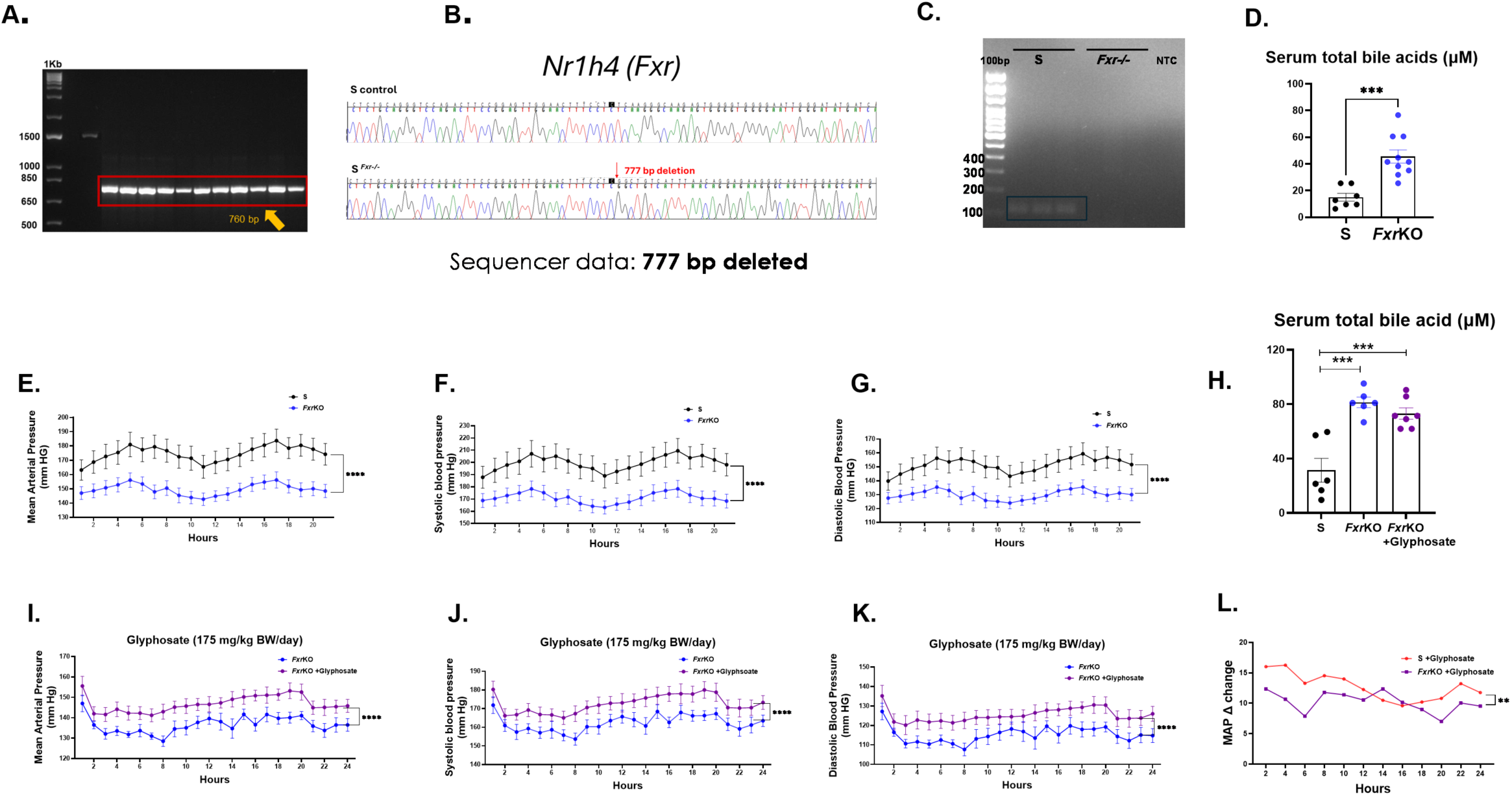
FXR Deletion Lowers Blood Pressure. **(A-C)** Screening and characterization of rats for CRISPR/Cas9-mediated deletion of the *Nr1h4 (Fxr)* locus. (**A)** Amplification of genomic DNA from S rat (1556 bp) and *Fxr*KO rats (760 bp). (**B)** Sequencer data confirmed a 777 bp deletion in *Fxr*KO rats within the *Nr1h4* locus. **(C)** Confirmation of exon 5 deletion in mRNA from the liver samples of S rat and *Fxr*KO rats by running the product of reverse transcription PCR on 2% agarose gel. **(D)** Comparison of serum total bile acids between control S and *Fxr*KO rats. n=7-10/groups. **(E-G)** Blood pressure comparisons between control S and *Fxr*KO rats: **(E)** Mean arterial pressure **(F)** systolic blood pressure, and **(G)** diastolic blood pressure (n = 7–10/group). **(H)** Comparison of the serum Total Bile Acids in S rats, *Fxr*KO rats and glyphosate-treatment *Fxr*KO.. (**I-K)** Blood pressure comparisons between *Fxr*KO rats with and without glyphosate treatment (175 mg/kg body weight daily): **(I)** Mean arterial pressure **(J)** Systolic blood pressure, and **(K)** Diastolic blood pressure (n = 6/group). **(L)** Comparison of the change in Mean Arterial pressure (ΔMAP) in S rat and *Fxr*KO rats after exposure to glyphosate. Data are presented as mean ± SEM. *p < 0.05, **p < 0.01, ***p < 0.001, ****p < 0.0001.

Disruption of FXR signaling was associated with a marked reduction in blood pressure, with significant decreases in MAP (149.7 mm Hg vs. 174.7 mm Hg, p<0.0001), systolic blood pressure (171.1 mm Hg vs. 199.6 mm Hg, p<0.0001) and diastolic blood pressure (130 mm Hg vs 151.4 mm Hg, p<0.0001) (**Figures 4E-G**). These findings serve as mechanistic evidence to demonstrate FXR signaling as a critical determinant of blood pressure regulation and that loss of FXR confers an anti-hypertensive phenotype.

Next, to address the important question of whether ablation of bile acid signaling via FXR was necessary to reverse the pro-hypertensive effect of glyphosate, *FxrKO* rats were exposed to glyphosate (175 mg/kg BW/day). Compared to glyphosate-treated wild-type rats, *Fxr*KO rats showed a substantially attenuated elevation in blood pressure. However, the bile acid levels remained comparable between *Fxr*KO rats treated with and without glyphosate exposure (**Figures 4H-K**). While these data demonstrate that glyphosate requires FXR signaling to mediate a prohypertensive effect, the partial attenuation of the hypertensive response in *Fxr*KO rats reveals that additional FXR-independent mechanisms are parallelly mediating the potent pro-hypertensive effect of glyphosate (**Figures 4L**).

### Microbial-derived Shikimate is a Pro-hypertensive Metabolite

To probe for FXR-signaling independent mechanisms, we focused on the microbial pathway inhibited by glyphosate. Glyphosate inhibits the shikimate pathway of microbiota, which can lead to accumulation of the substrate in the pathway, shikimic acid^10^. Accordingly, compared to control rats, shikimic acid levels were elevated in the rats exposed to glyphosate (**Figure 2E**). To test whether this elevated shikimic acid levels could contribute to increase in blood pressure, we examined the blood pressure effect of orally administered shikimic acid (100 mg/kg BW/day) in S rats. As shown in **Figures 5A-C**, rats exposed to shikimic acid showed elevated MAP (145.7 mmHg vs. 135.1 mmHg, p<0.0001), systolic blood pressure (177.9 mmHg vs. 164.8 mmHg, p<0.0001) and diastolic blood pressure (129.6 mmHg vs. 120.3 mmHg, p<0.0001). Further, similar to the rats exposed to glyphosate, the rats exposed to shikimic acid also demonstrated decreased vasorelaxation in responses to increasing concentrations of Ach (**Figure 5D-5H**) and had no effects on urinary protein excretion (**Figure 5I**). These data support our conclusion that as a result of glyphosate inhibiting the shikimate pathway, the substrate and microbial metabolite shikimate, accumulates in host circulation and elevates blood pressure.

**Figure 5.**
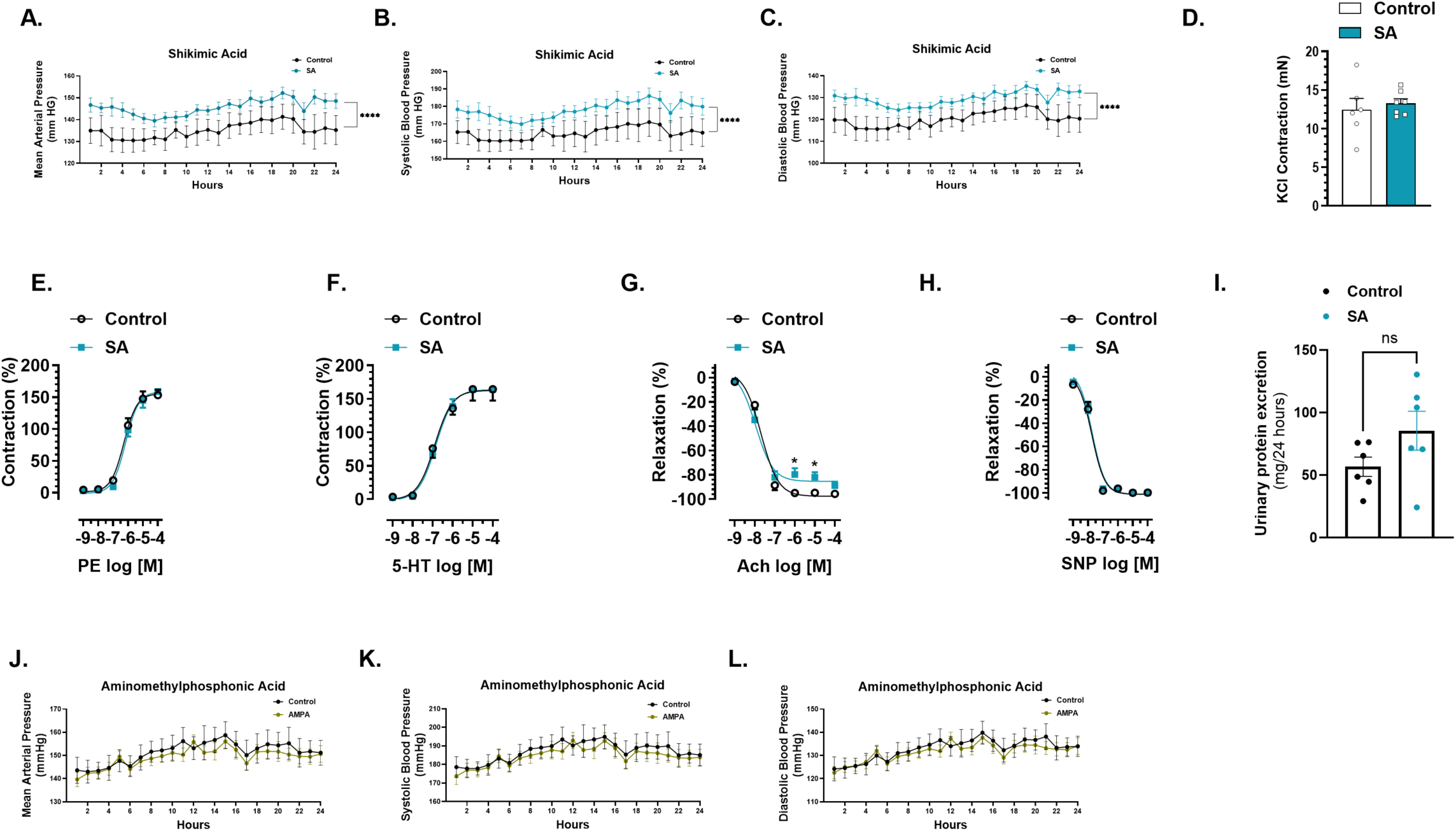
Shikimic Acid Administration, but not Aminomethylphosphonic Acid (AMPA) Increases Blood Pressure in S rats. **(A–C)** Blood pressure comparisons between control and shikimic acid-treated rats (100 mg/kg body weight daily): **(A)** Mean arterial pressure, **(B)** Systolic blood pressure, and **(C)** Diastolic blood pressure (n = 5-8/group). **(D)** KCl-induced contraction in mesenteric resistance arteries from control and shikimic acid-treated rats. **(E)** Contractions induced by phenylephrine (PE, 10⁻□–10^-4^ M). **(F)** Contractions induced by serotonin (5-HT, 10⁻□–10^-4^ M). **(G)** Endothelium-dependent relaxations induced by acetylcholine (ACh, 10⁻□–10^-4^ M). **(H)** Endothelium-independent relaxations induced by sodium nitroprusside (SNP, 10⁻□–10^-4^ M). Data are presented as mean ± SEM. ns = non-significant. **(I)** Urinary protein excretion comparisons between shikimic acid-treated and control rats. **(J–L)** Blood pressure comparisons between control and AMPA-treated rats (1.75 mg/kg body weight daily): **(J)** Mean arterial pressure. **(K)** Systolic blood pressure **(L)** Diastolic blood pressure rats (n = 6-9/group). Data are presented as mean ± SEM. ns = non-significant, *P < 0.05.

### The Endogenous Metabolite of Glyphosate, Aminomethylphosphonic Acid (AMPA), does not Lower Blood Pressure

In addition to the dual mechanisms of FXR-signaling and shikimate accumulation promoting hypertension, we considered a third factor, which is that glyphosate is metabolized into aminomethylphosphonic acid^22^. Therefore, we asked whether the increase in blood pressure in response to glyphosate exposure was due to its metabolite, aminomethylphosphonic acid (AMPA) because elevated levels of AMPA were also observed in serum and urine samples of rats exposed to glyphosate (**Figure 2C, D**). Unlike the rats exposed to glyphosate, rats orally administered with AMPA via drinking water did not increase their blood pressure compared to the control group (**Figure 5J-L**). These data support our conclusion that glyphosate, rather than its metabolite AMPA, is responsible for the observed increase in blood pressure.

## Discussion

Our laboratory was among the first to demonstrate that gut microbiota is linked to blood pressure regulation^15^. However, there is limited knowledge on the factors and mechanisms involved in gut microbiota remodeling that affect blood pressure. Environmental factors play a significant role in the pathogenesis of hypertension^23^. One such environmental factor is the use of agricultural herbicides. Glyphosate is the active ingredient in many agricultural and household weedicides, initially marketed under the trademark Roundup^1^. Interestingly, both the use of glyphosate and the prevalence of hypertension have been increasing. Human studies indicate that people are indeed exposed to glyphosate through dietary, occupational, and other environmental sources^24–26^. Moreover, there are several reports on the role of glyphosate in the pathophysiology of various diseases, such as reproductive and neurological disorders^27–30^, but, to our knowledge, none examining the hemodynamic effects of glyphosate. In this context, the current investigation is the first to establish a direct causal relationship between glyphosate exposure and hypertension. Using radiotelemetry, we observed rapid and sustained elevation in blood pressure in glyphosate-treated rats.

Next, our findings highlight the central role of gut microbiota in mediating glyphosate’s hypertensive effects. When glyphosate was administered subcutaneously, bypassing the gastrointestinal tract and direct interaction with gut microbiota, blood pressure remained unchanged. This strongly supports the notion that the adverse hemodynamic effect of glyphosate is mediated through its capacity to remodel gut microbiota rather than via direct systemic toxicity.

To further probe the mechanistic contribution of bile acid signaling in glyphosate-induced hypertension, we used a genetic approach targeting the FXR, a key regulator of bile acid homeostasis. Using the novel *Fxr*KO rats, we demonstrated that FXR signaling is required and is one of the factors mediating the prohypertensive function of glyphosate.

The role of bile acids has been studied in regulating lipid metabolism, glucose homeostasis, and vascular tone^33–36^. In addition, our previously published work demonstrates that the conjugated bile acid, taurocholic acid, is an antihypertensive bile acid^31^. We observed that taurocholic acid is elevated in *Fxr*KO rats compared to S rats (Supplementary Figure 3B). Administering glyphosate to *Fxr*KO rats decreased taurocholic acid (Supplementary Figure 4A, 4G). These data support the conclusion that the depletion of bile acid, and taurocholic acid in particular, may serve as a mechanistic link between gut microbiota and glyphosate-induced hypertension.

Further, *ex vivo* wire myography confirmed vascular impairment, that could be a potential additional contributor mediating glyphosate-induced hypertension. To investigate this further, we focused on the observed accumulation of shikimic acid levels, which occurs in response to glyphosate inhibiting the shikimate pathway of microbiota. Our data from rats exposed to shikimic acid showed vascular defects and elevated blood pressure, which serves as the second mechanistic insight.

One of the key findings in this study is the significant alteration in gut microbiota composition following glyphosate exposure that was accompanied by higher shikimic acid levels in circulation. Microbiome analysis revealed substantial reductions in alpha diversity, along with distinct shifts in bacterial community structure as shown by principal coordinate analysis. The shifts in gut microbiota composition at the species level, with significant reductions in commensal gut bacteria such as *Lactobacillus johnsonii*, *Bifidobacterium pseudolongum*, and *Faecalibacterium rodentium* are concordant with previous publications that these microbial taxa confer beneficial effects on cardiovascular health^32–36^. Beyond these specific commensals, our data support the conclusion by Mesnage *et al* that oral glyphosate exposure indeed remodeled gut microbiota composition^10^.

While the dosage used in our study (175 mg/kg BW/day) is below the observed-adverse-effect level (NOAEL), it is still higher than the Acceptable Daily Intake (ADI) of 1.75 mg/kg BW/day daily set by regulatory bodies such as the U.S. EPA. Higher doses have been effectively used in previous studies to uncover glyphosate’s biological effect^10,27,37^. However, for translational relevance, we conducted the low-dose glyphosate study and provided evidence for the first time to demonstrate that the currently acceptable daily intake of 1.75mg/kg BW/day also elevated blood pressure, albeit in a preclinical model.

In conclusion, our study commenced with 3 observations; (1) rising human exposure to an environmental agent, glyphosate, that can affect gut microbiota and (2) gut microbiota causally regulates blood pressure and (3) elevated blood pressure, a cardinal sign of health, is a global concern of modern society. By methodically addressing the question of whether these 3 factors are linked. At the time (1970’s) when glyphosate was released for agricultural practices, knowledge on the contributions of gut microbiota was limited, whereby, the effects of glyphosate in the shikimate pathway of microbiota was not in question. Now that microbiome science has conclusively proven its importance to our cardiovascular health, our study, which mechanistically addressed human associations, has revealed a provocative truth that glyphosate exposure leads to hypertension. This discovery holds significance because blood pressure is a key indicator of overall health, and elevated blood pressure is the primary risk factor for mortality among modern individuals, contributing to heart disease and stroke. Our discovery, albeit in an experimental rat model of hypertension, is alarming, and it prompts a re-evaluation of glyphosate toxicity in preclinical settings. Importantly, it sheds light on the potential widespread repercussions of the unintended consequences of the usage of glyphosate contributing to hypertension through its action on the gut microbiota, potentially affecting millions in the US alone.

## Methods

### Animals and Housing Conditions

The Institutional Animal Care and Use Committee of the University of Toledo College of Medicine and Life Sciences reviewed and approved all breeding and experimental procedures (reference numbers: 104573 for breeding and 108390 for experiments). All rats were maintained on a 12:12 hours light-dark cycle and water was always provided *ad libitum*. Rats were raised on a 0.3% NaCl containing diet (Harlan Teklad diet TD 7034, Madison, WI) until 8-10 weeks old, when they were switched to a 2% NaCl diet (Harlan Teklad diet TD94217, Madison, WI) and randomly assigned to experimental groups.

### BP Measurement Using Radiotelemetry

Adult 8-10 weeks old male S rats, an animal model that is genetically predisposed to hypertension, were used for the feeding studies^38^. Rats were surgically implanted with radiotelemetry transmitter probes and post recovery for 1 week, baseline blood pressure was measured as in our previous studies^15,39,40^. Groups one and two were administered with or without glyphosate (175 mg/kg BW/day) in their drinking water for 3 weeks. Groups three and four were administered with or without low dose glyphosate (1.75 mg/kg BW/day) in their drinking water for 3 weeks. Groups five and six were administered either with 1ml daily subcutaneous glyphosate (12 mg/ml) or sterile PBS for two weeks.

Next, two groups of 8–12-week-old male S rats were administrated with or without shikimic acid (100 mg/kg BW/day) for 5 weeks in their drinking water. Similarly, another two groups of 8-12-week-old male S rats were used for administration of AMPA (1.75 mg/kg BW/day) for 3 weeks.

### Generation of *Fxr* Knock Out (*Fxr*KO) Rat Model by CRISPR/CAS9 Gene Editing

Guide RNA was designed to target *Nr1h4* (*Fxr*) locus of intron 4 and downstream of intron 5 to remove exon 5. Oocyte microinjections were conducted at the University of Michigan Transgenic Animal Model core (Ann Arbor, MI). Genotyping was performed using the following primers: Forward (5’TTCGTAGGACCCTGTAGCATTACAAATTA3’) and Reverse (5’TCCCAAGTAAATACAGCCAAAGATCTAGC 3’). A potential founder rat was identified by the shorter size of DNA fragment than the non-founder. The identified founder rats were backcrossed to the Dahl Salt-Sensitive (S) rats, and their pups were intercrossed to obtain homozygosity of the disrupted *Fxr* allele. PCR products obtained from the homozygotes were shipped to Eurofins MWG Operon (https://www.eurofinsgenomics.com/en/home.aspx), for DNA sequencing (Sequencher 5.4.6.). To confirm FXR deletion at the mRNA levels, total RNA was extracted from liver tissues using the TRIzol method and reversed transcribed to generate cDNA^41^. The cDNA was used for PCR amplification with exon 5 specific primers: Forward 5’-TGGGAATGTTGGCTGAATGTTTG-3’ and Reverse 5’-CCCTTCGCTGTCCTCATTCA-3’. The relative expression of *Fxr* gene was quantified using the 2^−ΔΔCt^ method^42^.

Adult 8-10-week-old male *Fxr*KO and S rats were used for blood pressure comparisons using radiotelemetry. Similarly, another two groups of *Fxr*KO rats were administered with or without glyphosate (175 mg/kg BW/day) in their drinking water for 3 weeks and blood pressure was monitored using radiotelemetry.

### Chemicals

Glyphosate, N-(Phosphonomethyl) glycine (purity>95%) was obtained from Sigma-Aldrich (product number: 337757). Aminomethylphosphonic acid (AMPA) (99%) was obtained from Sigma-Aldrich (product number: 324817). Shikimic acid (purity≥99%) was obtained from Millipore Sigma (product number: S5375).

### 16S rRNA Gene Sequencing and Analysis of Microbiota Composition

#### 16S Polymerase Chain Reaction Library Preparation, Clean-Up, Normalization, and Pooling

Library preparation, clean-up, normalization, and pooling for 16S polymerase chain reaction were performed as described in our previous publications^39,40^. We followed the Illumina User Guide, 16S Metagenomic Sequencing Library Preparation—Preparing 16S Ribosomal RNA Gene Amplicons for the Illumina MiSeq System (Part No. 15044223 Rev. B). The V3-V4 region of the bacterial 16S rRNA gene was amplified by PCR using the Illumina sequencing primers: forward primer 5’ TCGTCGGCAGCGTCAGATGTGTATAAGAGACAGCCTACGGGNGGCWGCAG and reverse 5’TCTCGTGGGCTCGGAGATGTGTATAAGAGACAGGGACTACHVGGGTWTCTAAT. For index PCR, the Nextera XT index kit (FC-131-1002) from Illumina was used to attach dual indexes. Each 25 µL reaction mixture consisted of 2.5 µL of 10X reaction buffer (Invitrogen, Thermo Fisher Scientific), 0.5 µL of 10 mM dNTPs, 0.75 (for target PCR)/1 µL (for index PCR) of 50 mM MgCl2, 0.1 µL of 5U/µL of HotTaq polymerase (Invitrogen), 1 µL of each primer (5 µM) and 2.5 µL of 5 ng/µL DNA, with nuclease-free water added to reach a final volume of 25 µL. PCR amplification was performed in a BioRad T100TM thermal cycler (Hercules, CA). For the target amplification, the cycling conditions included: initial denaturation at 95°C for 5 min, followed by 25 cycles of 95°C for 30 s, 58°C for 30 s, 72°C for 30 s, with a final extension at 72°C for 5 min for target PCR. Index PCR was carried out in 8 cycles, with an initial denaturation at 95°C for 3 min, followed by 95°C for 30 s, 55°C for 30 s, 72°C for 30 s, and a final extension at 72°C for 5 min. Amplified products were purified twice using AMPure XP beads (Beckman Coulter Inc. Brea, CA). Concentrations of purified index PCR products was measured using the Qubit dsDNA HS Assay kit with Qubit 3.0 fluorometer (Life Technologies, Carlsbad, CA). Amplicons were pooled at equal molar concentrations of 4 nmol/L. The quality and size determination of the pooled library was performed using a 2100 Bioanalyzer (Agilent, Santa Clara, CA) before sequencing. Library Denaturing and MiSeq Sample Loading was done according to Illumina User Guide Illumina MiSeq System. The 10 pmol/L denatured and diluted library with 10% PhiX was loaded on an Illumina MiSeq V3 flow cell kit with 2×300 cycles.

#### Quality filtering, amplicon sequencing variant picking, data analysis

Chimeric sequences were identified and filtered using Quantitative Insights in Microbial Ecology version 2 (QIIME 2) (2021.11). The amplicon sequence variants (ASVs) were subsequently picked using QIIME 2, and taxonomy assignment was performed with ASVs using a pretained Naïve Bayes classifier utilizing Silva as the reference database clustered at 97%. Differentially abundant bacterial genera were analyzed by linear discriminant analysis (LDA) effect size (LEfSe) with LDA>2.

#### Nanopore GridION Metagenomics Sequencing of Fecal Samples

The SQK-LSK114 Ligation sequencing kit protocol (Nanopore) was followed.

#### DNA repair and end-prep

1 ug of DNA (47 μl volume) for each sample was mixed with1 μl DNA control strand, 3.5 μl NEBNext FFPE DNA repair buffer, 2 μl NEBNext FFPE DNA repair mix, 3.5 μl ultra II end-prep reaction buffer, and 3 μl ultra II end-prep enzyme mix. Using a thermal cycler, the DNA mix was incubated at 20°C for 5 mins followed by 65°C for 5 mins. Resuspended AMPure XP Beads (60 μl) (AXP) were added to the end-prep reaction mix in Eppendorf DNA LoBind tubes. After incubation on a rotator mixer for 5 mins at room temperature, magnetic rack was used to pellet the samples. The clear supernatant was discarded, and the pellet was washed twice with freshly prepared 200 ul of 80% ethanol. Then, the pellet without any residual ethanol was resuspended in 61 μl nuclease-free water. The resulting eluate (61 μl) was retained after pelleting out the beads on a magnet. Finally, 1 ul of eluted sample was quantified using Qubit fluorometer to measure the concentration of DNA.

#### Adapter ligation and clean-up

60 μl end-prepped DNA sample was mixed with 25 μl ligation buffer (LNB), 10 μl EBNext Quick T4 DNA ligase and 5 μl ligation adapter (LA) in a 1.5 ml Eppendorf LoBind tube. After incubating for 10 minutes at room temperature, 40 μl OF resuspended AMPure Beads were added to the samples. After incubation on a rotator mixer for 5 mins at room temperature, magnetic rack was used to pellet the samples. The supernatant was discarded, and the beads were washed twice with 250 μl Long Fragment Buffer (LFB). Then, the tubes were replaced back on the magnet to pellet the samples and resuspended in 15 μl elution buffer after pipetting off the supernatant. The resulting solution was incubated at 37°C and pelleted on the magnet to obtain clear and colorless eluate. The pelleted beads were discarded and the eluate containing the DNA library was retained. Qubit fluorometer was used to quantify DNA. The library was prepared to 12 μl at 10-20 fmol.

#### Loading prepared library onto the SpotON flow cell

The prepared library was used for loading into the flow cell. 12 ul of the DNA library was mixed with 37.5 ul of Sequencing buffer and 25.5 ul of Library solution and loaded on to SpotON flow cell (10.4.1) separately. The preparation of the priming mix and loading of library onto the GridION flow cell was performed as instructed by the Ligation Sequencing Kit V14 (SQK-LSK114) protocol. The Flowcells (10.4.1) were allowed to run for 72 hours on A GridION with high accuracy base calling.

### Data analysis

The Fastq files (passed) were exported and analyzed using the CZ-ID (Chan Zuckerberg initiative)^43^ portal after filtering of host and human reads followed by metagenomic pipeline. The normalized 1 million bases of sequenced reads (of Taxa at species level) were further analyzed on Microbiome Analyst to visualize the differential abundance of species.

### Total Bile Acid Quantification

Serum total bile acid levels were measured using the Diazyme Total Bile Acid (TBA) assay kit (DZ042A-KY1) according to the manufacturer’s protocol.

### Bile Acid Quantitation by Ultra-High Performance Liquid Chromatography-Tandem Mass Spectrometry

The experimental methodology used to quantify bile acid levels in serum was based on a previous publication with minor modifications^44^. Targeted quantification of bile acids in serum was determined using a Vanquish UHPLC system, a TSQ Quantis Triple Quadrupole mass spectrometer (Thermo Fisher Scientific), and an ACQUITY C8 BEH UPLC column (2.1 × 100 mm, 1.7 μm) (Waters). Protein precipitation was achieved by mixing 50 μL serum with 150 μL ice-cold methanol and 0.5 μmol/L deuterated internal standards. After 20 minutes of incubation at −20°C, the samples were centrifuged at 12,000 g for 10 mins at 4°C, and 150 μL of supernatants were transferred to autosampler vials. Bile acids were determined using various reaction monitoring and ion monitoring modalities. Finally, the results were quantified using Skyline (MacCoss Lab Software), which compared integrated peak areas against standard curves.

### Vascular Function

*Ex vivo* wire myography was performed as described^45^. Immediately after euthanasia, thoracic aorta and third- to fourth-order mesenteric resistance arteries (MRAs) were dissected and gently cleared of surrounding connective and adipose tissue in cold, oxygenated modified Krebs-Henseleit buffer (130 mM NaCl, 4.7 mM KCl, 1.17 mM MgSO₄, 1.18 mM KH₂PO₄, 1.6 mM CaCl₂, 25 mM NaHCO₃, 5.5 mM glucose, and 0.03 mM EDTA; pH 7.4). The buffer was continuously aerated with a 95% O₂ and 5% CO₂ gas mixture throughout the experiment.

Aortic rings and MRAs (∼2 mm in length) were mounted on a DMT 620M multi-channel myograph system, with aortas placed in pin myographs and MRAs in wire myographs. Tissues were maintained in 37°C oxygenated Krebs buffer. After stabilization, aortic rings were incrementally stretched to a resting tension of 10 mN. MRAs were normalized using the manufacturer’s protocol to approximate physiological stretch, employing a target pressure of 13.3 kPa and an internal circumference ratio (IC₁/IC₁₀₀) of 0.9. A stable pretension was achieved followed by multiple wash steps of the vessels with fresh 37°C oxygenated Krebs buffer for 45 mins.

To assess vascular function, isolated aortic rings or MRAs with intact endothelium were challenged with a high-KCL solution (120 mM), vasoconstrictors PE or serotonin (5-HT) (10^−9^–10^−4^ M), endothelial-dependent vasodilator, Ach (10^−9^–10^−4^ M), or an endothelial-independent vasodilator, sodium nitroprusside (SNP) (10^−9^–10^−5^ M). Vessels were thoroughly washed between treatments with 37°C Krebs buffer until baseline tone was re-established.

### Urinary Protein Excretion (UPE)

UPE was quantified as previously published^40^. Briefly, each rat was housed individually in a metabolic cage and urine samples were collected over a 24-hr period. The pyrogallol based QuanTtest Red Total Protein Assay from Quantimetrix (Redondo Beach, CA, USA) was used to measure protein concentrations of the urine samples.

### Targeted Quantification of Glyphosate and Aminomethylphosphonic acid (AMPA)

Certified standard solutions of glyphosate, AMPA, ^13^C_2_,^15^N-glyphosate, and ^13^C,^15^N,D_2_-AMPA (in water) with concentrations at 100 µg/mL and purities ≥ 95% were purchased from Cambridge Isotope Laboratories (Andover, MA, USA). Oasis MCX (60 mg/3 mL) and MAX (60 mg/3 mL) solid-phase extraction (SPE) cartridges were purchased from Waters Corporation (Milford, MA, USA). Methanol (MeOH), acetonitrile (ACN), water, formic acid (88%), and ammonium hydroxide (NH_4_OH; 28–30%) of LC-MS grade were obtained from Fisher Scientific (Waltham, MA, USA). Methylenediphosphonic acid (10 mM) was obtained from Restek Corporation (Bellefonte, PA, USA).

Glyphosate and AMPA were extracted from urine and serum using a procedure described previously with minor modifications^46^. Briefly, a 250 µL aliquot of each urine or serum sample was transferred into a 15-mL polypropylene (PP) tube. Then, 2.5 ng of internal standards were spiked, 250 µL of 100 µM methylenediphosphonic acid was added, and the sample was allowed to equilibrate at room temperature for 30 min. The sample was then loaded onto an Oasis MCX (60 mg/3 mL) cartridge that had been sequentially preconditioned with 1.5 mL MeOH and 1.5 mL water. The cartridge was washed with 2 mL water, and the elutes were collected immediately. Thereafter, 3 mL of 3% NH_4_OH (v/v) was added, and the mixture was vortexed vigorously. The sample was then loaded onto an Oasis MAX cartridge (60 mg/3 mL) that had been sequentially preconditioned with 1.5 mL MeOH, 1.5 mL water, and 1.5 mL of 3% NH_4_OH. The cartridge was washed with 2 mL of 3% NH_4_OH and 2 mL MeOH, followed by vacuum-drying for 1 min. The analytes were then eluted into a 15-mL PP tube using MeOH containing 3% formic acid, and the elute was evaporated to near dryness under a nitrogen stream at 36°C. The residue was reconstituted in 250 µL ACN:water (5:95, v/v) containing 0.1% formic acid and 2.5 µM methylenediphosphonic acid, which was injected into liquid chromatography-tandem mass spectrometry (LC-MS/MS) for analysis.

Identification and quantification of glyphosate and AMPA were performed using an ABSciex 5500+ Q-trap mass spectrometer (Framingham, MA, USA) interfaced with an ExionLC ultra-high performance liquid chromatography (UPLC) (Sciex; Redwood City, CA, USA). Chromatographic separation of the analytes was performed on a Germini^®^ C6-Phenyl column (150 × 4.6 mm, 5 µm; Phenomenex, Torrance, CA, USA) connected to a Betasil C18 guard column (20 × 2.1 mm, 5 µm; Thermo Fisher Scientific, Waltham, MA, USA). The mobile phases, maintained at a flow rate of 0.6 mL/min, were (A) water containing 0.1% formic acid (v/v) and 2.5 µM LC passivation solution (methylenediphosphonic acid) and (B) acetonitrile containing 0.1% formic acid (v/v). The following gradient program was applied: hold at 5% B for 2 min, linear ramp to 95% B over 8 min, hold at 95% B for 1 min, then return to initial conditions in over 1 min, and equilibrate for another 2 min prior to the next injection. The column temperature was maintained at 40°C.

The analytes were determined using multiple reaction monitoring (MRM) negative-ion mode. The IonSpray voltage was −4500 V; the ionization source temperature was 500°C; and the curtain gas flow was 20 psi.

### Untargeted Quantification of Primary Metabolites

Serum metabolite profiling was performed at the University of California, Davis West Coast Metabolomics Center. Serum samples were used for extraction using 1mL of a 3:3:2 (v/v/v) mixture of acetonitrile:isopropanol:water (ACN:IPA:H2O). Half of each extract was dried completely and derivatized by adding 10 µL of 40 mg/mL of methoxyamine in pyridine, followed by shaking at 30 °C for 1.5 hours. Subsequently, 91 µL of N-Methyl-N-(trimethylsilyl) trifluoroacetamide (MSTFA) and Fatty Acid Methyl Esters (FAMEs) mixture was added to each sample, and the mixture were shaken at 37 °C for 30 minutes to complete derivatization. Samples were then vialed, capped, and injected onto the instrument.

Analysis was conducted using an Agilent 7890A gas chromatograph (GC) coupled with a LECO time-of-flight mass spectrometer (TOF-MS). A 0.5□µL aliquot of each derivatized sample was injected in splitless mode onto a RESTEK RTX-5Sil MS column with an Integra-Guard pre-column at an inlet temperature of 275□°C and helium as the carrier gas at a flow rate of 1□mL/min. The GC oven temperature was programmed to hold at 50□°C for 1 minute, then ramp at 20□°C/min to 330□°C, where it was held for 5 minutes. The transfer line and electron impact (EI) ion source were set to 280□°C and 250□°C, respectively. Mass spectra were acquired over the m/z range of 85–500 at a rate of 17 spectra per second.

## Supplementary Material

Figures S1-S4

## Data Availability

All data and materials have been included within the manuscript. Whole genome sequence data is reposited in the SRA database with BioProject number **PRJNA1418581**. 16S rRNA sequencing data is reposited in the SRA with BioProject number **PRJNA1416381**.

## Statistical Analyses

All statistical analyses were performed using GraphPad Prism version 10.3.1 for Windows (GraphPad Software, Boston, MA, USA; www.graphpad.com). Differences between two groups were assessed using an unpaired, two-tailed Student’s *t*-test with a 95% confidence interval. Twenty-four-hour blood pressure and *ex vivo* vascular wire myography measurements were analyzed using two-way ANOVA followed by Fisher’s least significant difference (LSD) test. Correlation analyses were performed using Pearson’s correlation coefficient, assuming a Gaussian distribution of the data. Data are presented as mean ± SEM, and statistical significance is reported as *p-values*.

---

**Supplementary Figure 1:**
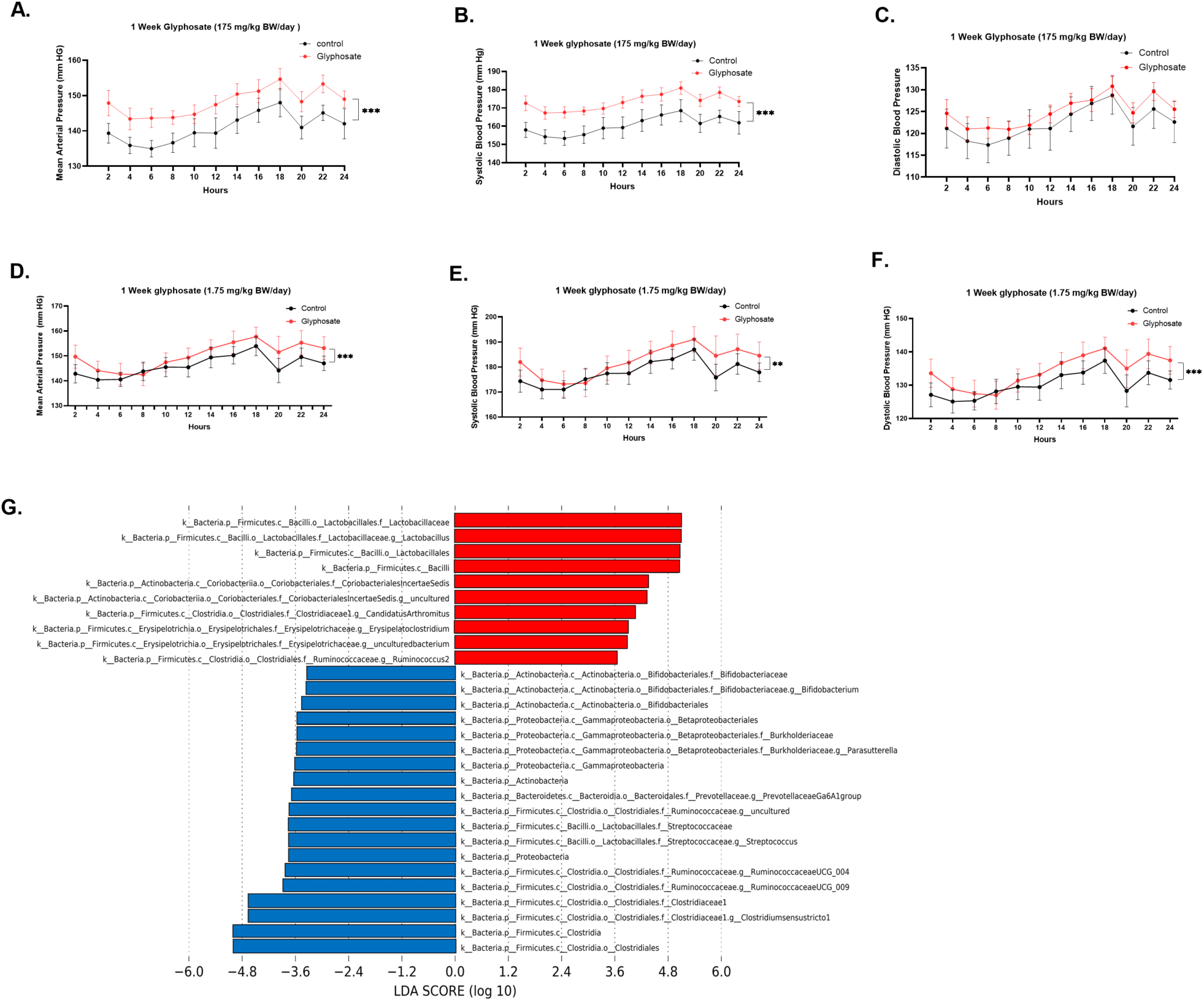
**(A–C)** Blood pressure comparisons between control and glyphosate-treated rats (175 mg/kg body weight daily) at week 1: **(A)** Mean arterial pressure **(B)** Systolic blood pressure **(C)** Diastolic blood pressure (n = 7–8/group). (D-F) Blood pressure comparisons between control and glyphosate-treated rats (1.75 mg/kg body weight daily) at week 1: **(D)** Mean arterial pressure **(E)** Systolic blood pressure **(F)** Diastolic blood pressure (n = 7–8/group).**(G)** 16S rRNA gene sequencing followed by Linear Discriminant Analysis Effect Size (LEfSe) identified differentially abundant bacterial taxa (LDA score > 2.0, P<0.05) between control and glyphosate-treated rats. Bar plots (red indicates the control group, blue indicate *Fxr*KO group) display enriched taxa in each group.

**Supplementary Figure 2:**
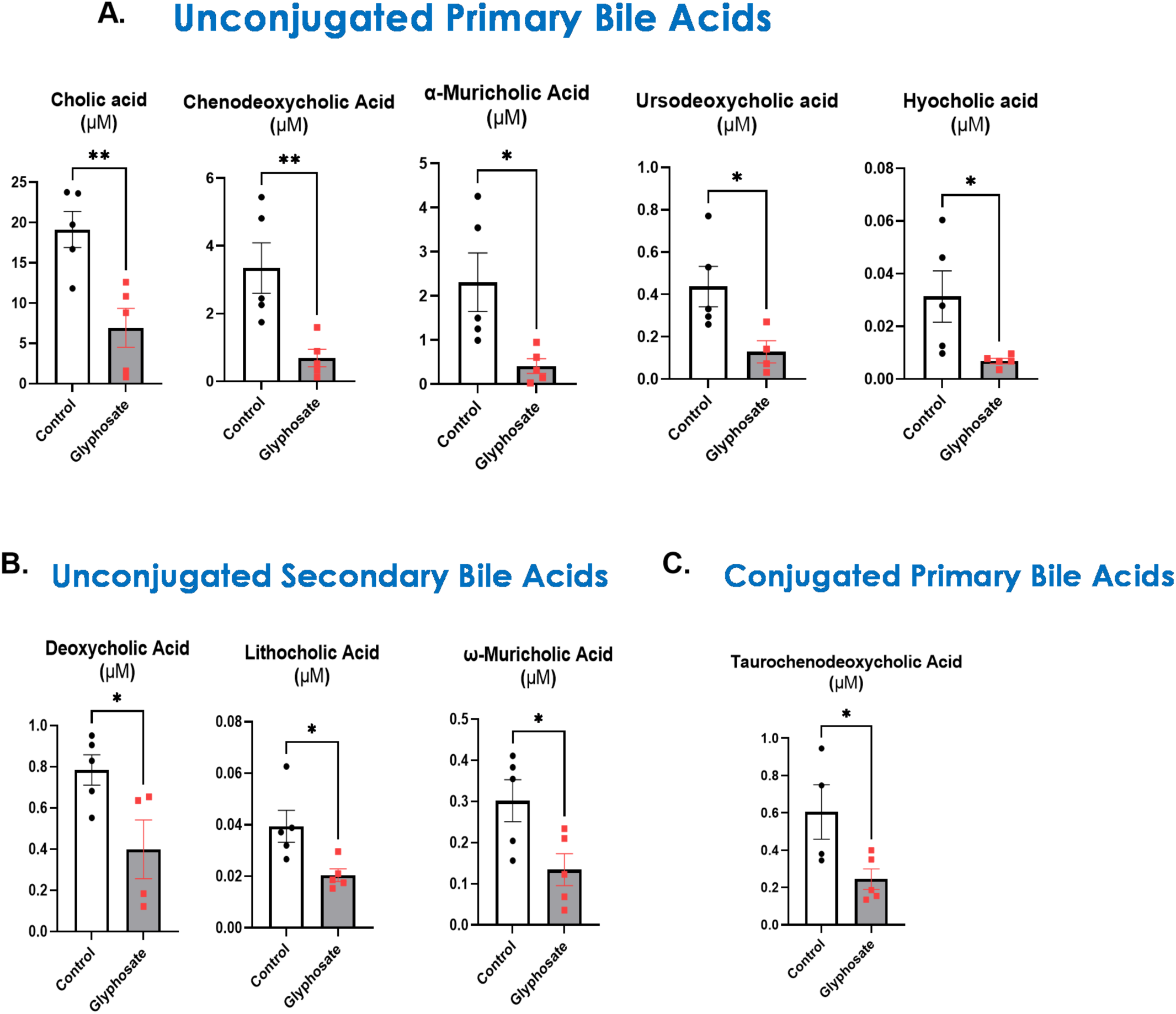
Glyphosate Lowered Circulating Bile Acids. **(A)** Unconjugated primary bile acids. **(B)** Unconjugated secondary bile acids. (TBA). (**C)** Unconjugated secondary bile acids. n=5/group. ns = non-significant, *p<0.05, **p<0.01.

**Supplementary Figure 3.**
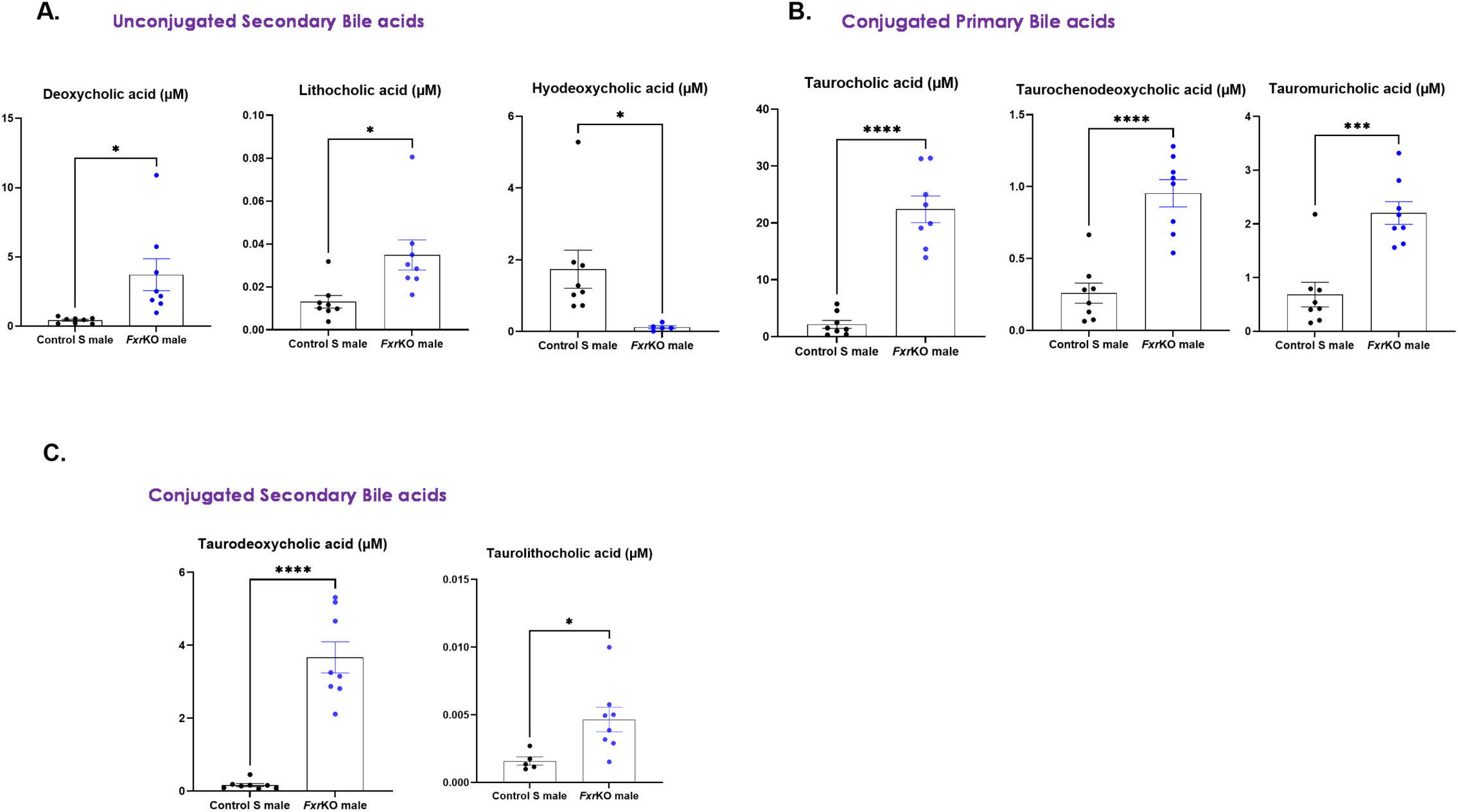
*Fxr*KO Showed Higher Levels of Circulating Bile Acids than S rats. Targeted bile acid quantification in sera of S rat and *Fxr*KO rats: **(A)** Unconjugated secondary bile acids. **(B)** Conjugated primary bile acids. **(C)** Conjugated secondary bile acids. n=8/group. ns = non-significant, *p<0.05, **p<0.01, ***p<0.001, ****p<0.0001.

**Supplementary Figure 4:**
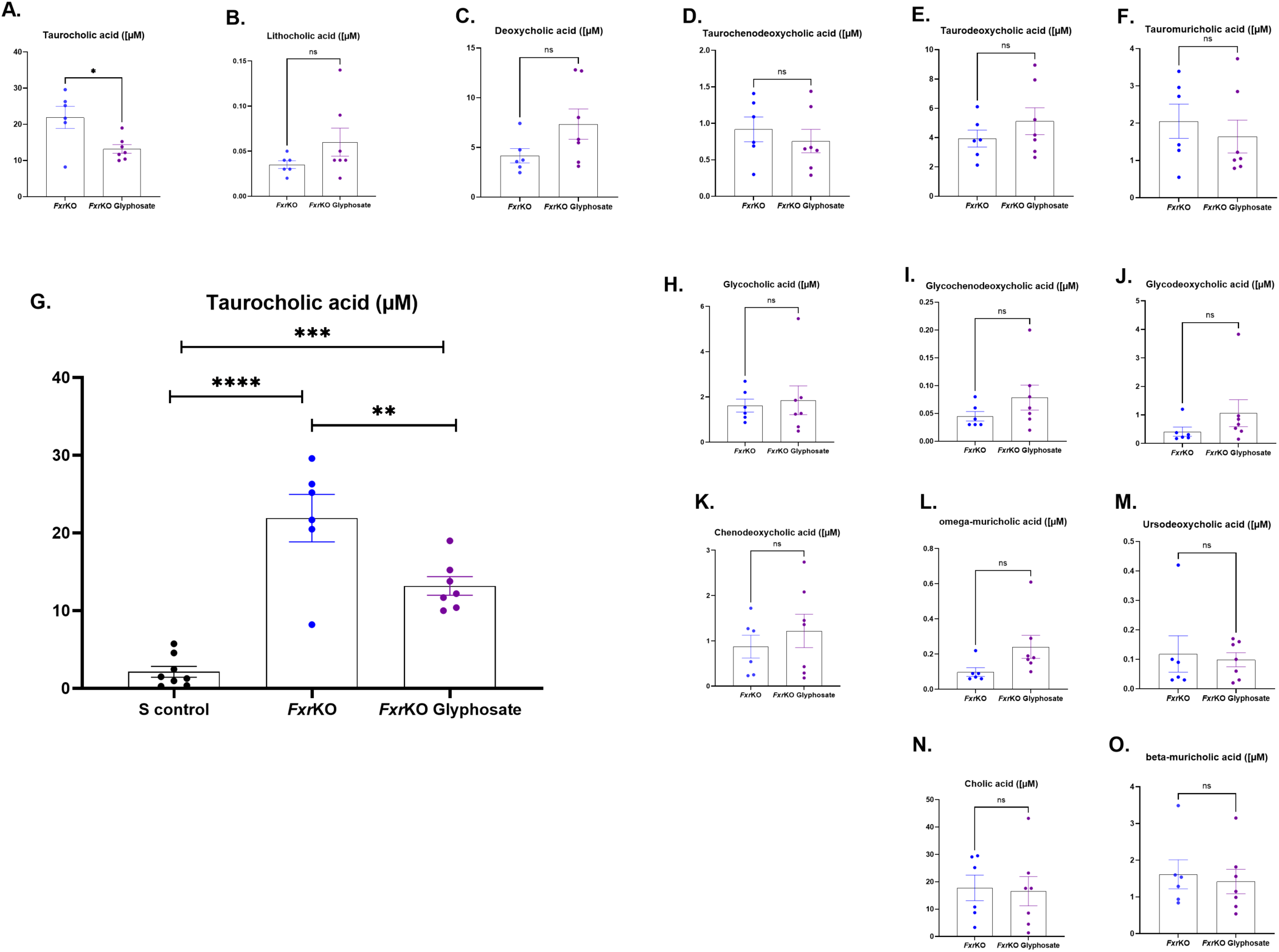
Targeted bile acid quantification in sera. n=6-8/group. ns = non-significant, *p<0.05, **p<0.01, ***p<0.001, ****p<0.0001.

## References

1. Mendelson, J. Roundup: the world’s biggest-selling herbicide. Ecologist 28, 254 (1998).

2. Benbrook, C. M. Trends in glyphosate herbicide use in the United States and globally. Environ. Sci. Eur. 28, 1–15 (2016).

3. Duke, S. O. & Powles, S. B. Glyphosate: a once-in-a-century herbicide. Pest Manag. Sci. Former. Pestic. Sci. 64, 319–325 (2008).

4. Guyton, K. Z. et al. Carcinogenicity of tetrachlorvinphos, parathion, malathion, diazinon, and glyphosate. Lancet Oncol. 16, 490–491 (2015).

5. Li, Z.-M., Jeong, H. & Kannan, K. Occurrence and profiles of glyphosate and aminomethylphosphonic acid in paired urine and feces of humans, cats, and dogs. Environ. Int. 109497 (2025).

6. Ospina, M. et al. Exposure to glyphosate in the United States: data from the 2013–2014 National Health and Nutrition Examination Survey. Environ. Int. 170, 107620 (2022).

7. Jaeger, B. C. et al. Hypertension statistics for US adults: an open-source web application for analysis and visualization of national health and nutrition examination survey data. Hypertension 80, 1311–1320 (2023).

8. Schönbrunn, E. et al. Interaction of the herbicide glyphosate with its target enzyme 5-enolpyruvylshikimate 3-phosphate synthase in atomic detail. Proc. Natl. Acad. Sci. 98, 1376–1380 (2001).

9. Knaggs, A. R. The biosynthesis of shikimate metabolites. Nat. Prod. Rep. 20, 119–136 (2003).

10. Mesnage, R. et al. Use of shotgun metagenomics and metabolomics to evaluate the impact of glyphosate or Roundup MON 52276 on the gut microbiota and serum metabolome of Sprague-Dawley rats. Environ. Health Perspect. 129, 17005 (2021).

11. Structure, function and diversity of the healthy human microbiome. Nature 486, 207–214 (2012).

12. Turnbaugh, P. J. et al. The human microbiome project. Nature 449, 804–810 (2007).

13. Yang, T. et al. Gut dysbiosis is linked to hypertension. Hypertension 65, 1331–1340 (2015).

14. Joe, B. et al. Microbiota Introduced to Germ-Free Rats Restores Vascular Contractility and Blood Pressure. Hypertension 76, 1847–1855 (2020).

15. Mell, B. et al. Evidence for a link between gut microbiota and hypertension in the Dahl rat. Physiol. Genomics 47, 187–197 (2015).

16. Galla, S. et al. Disparate effects of antibiotics on hypertension. Physiol. Genomics 50, 837–845 (2018).

17. Santisteban, M. M. et al. Hypertension-linked pathophysiological alterations in the gut. Circ. Res. 120, 312–323 (2017).

18. Adnan, S. et al. Alterations in the gut microbiota can elicit hypertension in rats. Physiol. Genomics 49, 96–104 (2017).

19. Sherman, S. B. et al. Prenatal androgen exposure causes hypertension and gut microbiota dysbiosis. Gut Microbes 9, 400–421 (2018).

20. Bardhan, P. et al. Salt-Responsive Gut Microbiota Induces Sex-Specific Blood Pressure Changes. Circ. Res. 135, 1122–1137 (2024).

21. Joe, B. Dr Lewis Kitchener Dahl, the Dahl rats, and the “inconvenient truth” about the genetics of hypertension. Hypertension 65, 963–969 (2015).

22. Anadón, A. et al. Toxicokinetics of glyphosate and its metabolite aminomethyl phosphonic acid in rats. Toxicol. Lett. 190, 91–95 (2009).

23. Sharma, P. & Brook, R. D. Echoes from Gaea, Poseidon, Hephaestus, and Prometheus: environmental risk factors for high blood pressure. J. Hum. Hypertens. 32, 594–607 (2018).

24. Gillezeau, C. et al. The evidence of human exposure to glyphosate: a review. Environ. Heal. 18, 1–14 (2019).

25. Muñoz, J. P., Silva-Pavez, E., Carrillo-Beltrán, D. & Calaf, G. M. Occurrence and exposure assessment of glyphosate in the environment and its impact on human beings. Environ. Res. 231, 116201 (2023).

26. Li, Z.-M., Jeong, H. & Kannan, K. Widespread occurrence of glyphosate and aminomethylphosphonic acid in indoor dust from urban homes across the United States and its contribution to human exposure. Environ. Int. 192, 109005 (2024).

27. Winstone, J. K. et al. Glyphosate infiltrates the brain and increases pro-inflammatory cytokine TNFα: implications for neurodegenerative disorders. J. Neuroinflammation 19, 193 (2022).

28. Liu, J.-B., Li, Z.-F., Lu, L., Wang, Z.-Y. & Wang, L. Glyphosate damages blood-testis barrier via NOX1-triggered oxidative stress in rats: Long-term exposure as a potential risk for male reproductive health. Environ. Int. 159, 107038 (2022).

29. Hsiao, Y.-C. et al. Evaluation of neurological behavior alterations and metabolic changes in mice under chronic glyphosate exposure. Arch. Toxicol. 98, 277–288 (2024).

30. Mendez, F., Ordoñez-Betancourth, J. & Abrahams, N. Effects of glyphosate exposure on reproductive health: A systematic review of human, animal and in-vitro studies. Expo. Heal. 14, 635–669 (2022).

31. Chakraborty, S. et al. Conjugated bile acids are nutritionally re-programmable antihypertensive metabolites. J. Hypertens. 41, 979 (2023).

32. Krautkramer, K. A., Fan, J. & Bäckhed, F. Gut microbial metabolites as multi-kingdom intermediates. Nat. Rev. Microbiol. 19, 77–94 (2021).

33. Jie, Z. et al. The gut microbiome in atherosclerotic cardiovascular disease. Nat. Commun. 8, 845 (2017).

34. Xiong, Y. et al. Lactobacillus induced by irbesartan on spontaneously hypertensive rat contribute to its antihypertensive effect. J. Hypertens. 42, 460–470 (2024).

35. Miao, H. et al. Targeting Lactobacillus johnsonii to reverse chronic kidney disease. Signal Transduct. Target. Ther. 9, 195 (2024).

36. Wang, J. et al. Clostridium butyricum and Bifidobacterium pseudolongum attenuate the development of cardiac fibrosis in mice. Microbiol. Spectr. 10, e02524–22 (2022).

37. Bartholomew, S. K. et al. Glyphosate exposure exacerbates neuroinflammation and Alzheimer’s disease-like pathology despite a 6-month recovery period in mice. J. Neuroinflammation 21, 1–19 (2024).

38. Rapp, J. P. & Dene, H. Development and characteristics of inbred strains of Dahl salt-sensitive and salt-resistant rats. Hypertension 7, 340–349 (1985).

39. Mei, X. et al. Repurposing a Drug Targeting Inflammatory Bowel Disease for Lowering Hypertension. J. Am. Heart Assoc. 11, e027893 (2022).

40. Chakraborty, S. et al. Salt-responsive metabolite, β-hydroxybutyrate, attenuates hypertension. Cell Rep. 25, 677–689 (2018).

41. Rio, D. C., Ares, M. J., Hannon, G. J. & Nilsen, T. W. Purification of RNA using TRIzol (TRI reagent). Cold Spring Harb. Protoc. 2010, pdb.prot5439 (2010).

42. Schmittgen, T. D. & Livak, K. J. Analyzing real-time PCR data by the comparative C(T) method. Nat. Protoc. 3, 1101–1108 (2008).

43. Simmonds, S. E. et al. CZ ID: a cloud-based, no-code platform enabling advanced long read metagenomic analysis. bioRxiv 2002–2024 (2024).

44. Tian, Y., Cai, J., Allman, E. L., Smith, P. B. & Patterson, A. D. Quantitative analysis of bile acid with UHPLC-MS/MS. Transl. Bioinforma. Ther. Dev. 291–300 (2021).

45. Grano de Oro, A., et al. Spontaneous vascular dysfunction in Dahl salt-sensitive male rats raised without a high-salt diet. Physiol. Rep. 12, e16165 (2024).

46. Li, Z.-M. & Kannan, K. A method for the analysis of glyphosate, aminomethylphosphonic acid, and glufosinate in human urine using liquid chromatography-tandem mass spectrometry. Int. J. Environ. Res. Public Health 19, 4966 (2022).

